# Structural analysis of S-ring composed of FliFG fusion proteins in marine *Vibrio* polar flagellar motors

**DOI:** 10.1101/2024.04.26.591406

**Authors:** Norihiro Takekawa, Tatsuro Nishikino, Jun-ichi Kishikawa, Mika Hirose, Miki Kinoshita, Seiji Kojima, Tohru Minamino, Takayuki Uchihashi, Takayuki Kato, Katsumi Imada, Michio Homma

## Abstract

The marine bacterium *Vibrio alginolyticus* possesses a polar flagellum driven by a sodium ion flow. The main components of the flagellar motor are the stator and rotor. The C-ring and MS-ring which are composed of FliG and FliF, respectively, are parts of the rotor. Here, we purified an MS-ring composed of FliF-FliG fusion proteins and solved the near-atomic resolution structure of the S-ring—the upper part of the MS-ring—using cryo-electron microscopy. This is the first report of an S-ring structure from *Vibrio* whereas, previously, only those from *Salmonella* have been reported. The *Vibrio* S-ring structure reveals novel features compared to that of *Salmonella* such as tilt angle differences of the core domain and the β-collar region, the decrease of the inter-subunit interaction between core domains, and altered electrostatic inner-surface. The residues potentially interact with other flagellar components, such as FliE and FlgB, are well structurally conserved in *Vibrio* S-ring. These comparisons clarified the conserved and non-conserved structural features of the MS-ring across different species.

**IMPORTANCE:** Understanding the structure and function of the flagellar motor in bacterial species is essential for uncovering the mechanisms underlying bacterial motility and pathogenesis. Our study revealed the structure of the *Vibrio* S-ring, a part of its polar flagellar motor, and highlighted its unique features compared with the well-studied *Salmonella* S-ring. The observed differences in the inter-subunit interactions and in the tilt angles between the *Vibrio* and *Salmonella* S-rings highlighted the species-specific variations in the flagellar assembly. By concentrating on the region where the S-ring and the rod proteins interact, we uncovered conserved residues essential for the interaction. Our research contributes to advancing of bacterial flagellar biology.

## INTRODUCTION

The marine bacterium *Vibrio alginolyticus—*hereinafter referred to as “*Vibrio*”— has two types of flagella: a polar flagellum for swimming in an aqueous environment and lateral flagella for swarming on the surface (1) (Fig. 1A). The polar and lateral flagella are driven by sodium ion (Na^+^) and proton (H^+^) flow, respectively. The flagellar motor is a force-generating complex at the flagellum base, consisting of a dozen stator units and a rotor for rotating the flagellar filament, which acts as a helical propeller. The stator is an energy conversion unit composed of two types of membrane proteins: MotA and MotB for the H^+^-driven flagellum and PomA and PomB for the Na^+^-driven flagellum (2).

**FIG 1.**
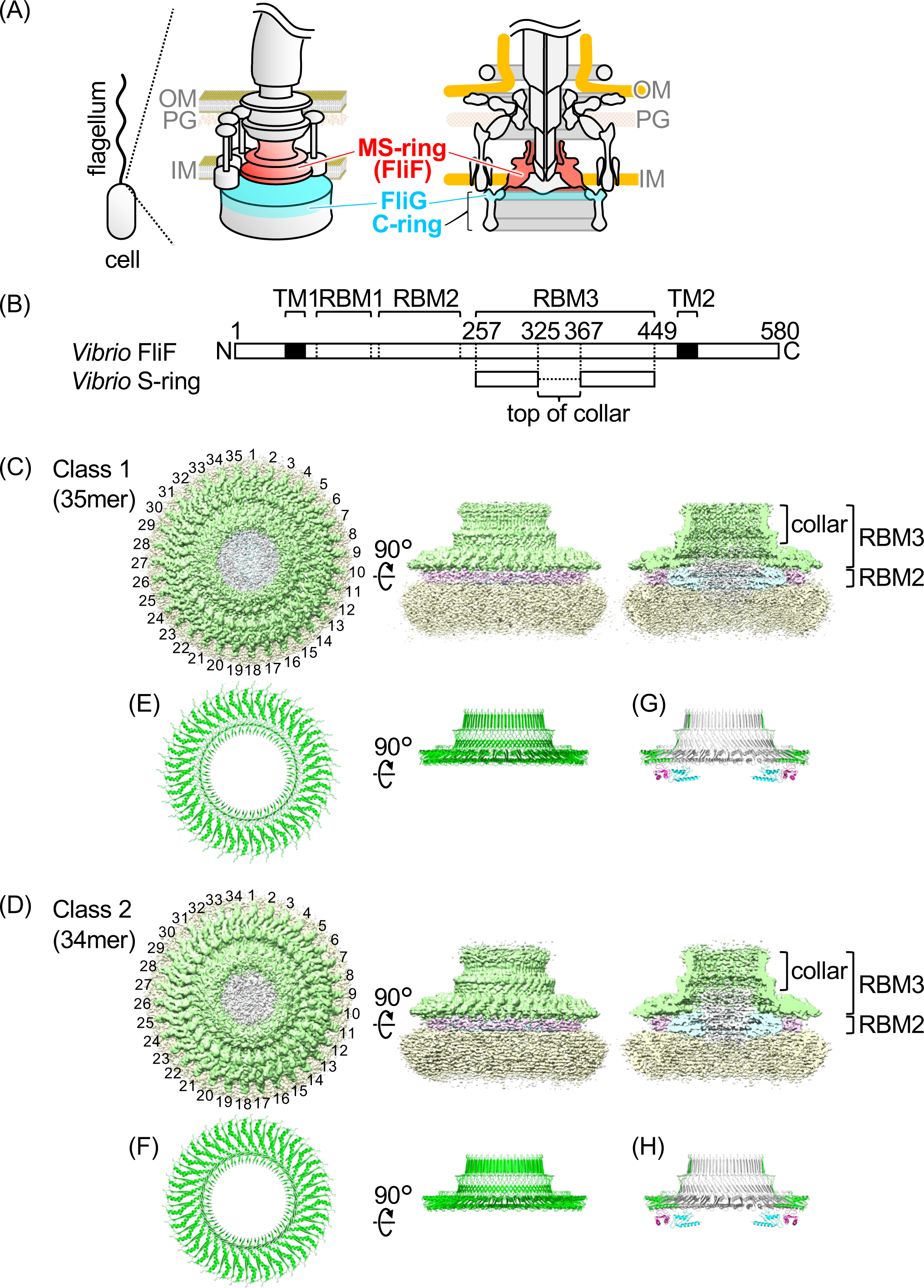
Flagellar motor structure and MS-ring (FliF) in *Vibrio*. (A) Schematic diagram of the flagellar basal body in *Vibrio*. The *Vibrio* cell has a single flagellum at its cell pole and there is a flagellar motor at the flagellar base. The MS-ring— consisting of FliF—and the upper part of the C-ring—consisting of FliG—are colored with red and cyan, respectively. OM: outer membrane, PG: peptidoglycan layer, IM: inner membrane. (B) Schematic representation of the FliF primary structure containing the region forming the RBM1, RBM2, RBM3 (S-ring), and transmembrane helices (TM1 and TM2) with residue numbers. (C and D) Surface representation and vertical section of the cryo-EM map of the purified MS-ring without rotational symmetry correction (C1) of class 1 (35-mer) and class 2 (34-mer). Green: S-ring, cyan: inner part of the M-ring, pink: middle part of the M-ring, yellow: unstructured outermost region of the M-ring, gray: unstructured innermost region. (E and F) The Cα ribbon drawings of the S-ring atomic models built from the maps after rotational symmetry correction (C35 for Class 1 and C34 for Class 2). (G and H) The Cα ribbon drawings of the S-ring atomic models with the previously reported structures of RBM2 (8).

The stator interacts with the rotor—composed of a C-ring and an MS-ring—to generate torque through a gear-like motion (3). The C-ring consists of three soluble proteins: FliG, FliM, and FliN (4, 5) and is composed of ∼34, ∼34, and ∼102 molecules of FliG, FliM, and FliN, respectively (6, 7). MotA or PomA in the stator interact with FliG in the rotor. The C-ring is tightly attached to the MS-ring, which consists of ∼34 FliF molecules with two transmembrane helices (8, 9). FliF contains three ring-building motifs (RBMs): RBM1, RBM2, and RBM3 in the periplasmic region between the two transmembrane helices (Fig. 1B). Among these motifs, RBM3 forms an S-ring on the distal side of the cytoplasm within the MS-ring.

Flagellar construction is postulated to start with the formation of the transmembrane export gate complex and MS ring, which work as the foundation for flagellar formation (10–12). FliF from *Salmonella enterica* serovar Typhimurium—hereinafter referred to as “*Salmonella*”—efficiently forms the MS-ring by its overproduction in *E. coli* (13). In this case, the 32–36 RBM3 subunits form an S-ring, whereas the 21–23 RBM2 subunits form an inner M-ring at the bottom of the S-ring. In addition, the MS-ring, attached to the C-ring, was isolated by the overproduction of *Salmonella* FliF/FliG/FliM/FliN proteins in *E. coli* (14). Using this system, it was tried to solve the MS-ring and C-ring structures using cryo-electron microscopy (cryo-EM); however, only RBM3 and RBM2 of the MS-ring were solved (15). The MS-ring contains 32–34 FliF subunits, with 33 being the most common (15) while 34 RBM3s form the S-ring, and 23 RBM2s form the inner part of the M-ring in the intact basal body containing the export gate complex and rod in *Salmonella* (9).

*Vibrio* FliF rarely forms the MS-ring by overproduction in *E. coli* but forms unstructured soluble protein complexes (16)*. Vibrio* FliF requires FlhF—an essential factor in the generation of the single polar flagellum in *Vibrio—*to determine the flagellar number at the cell pole along with FlhG or FliG for efficient MS-ring formation in *E. coli*; in other words, *Vibrio* FliF efficiently forms the MS-ring by co-expression with FlhF or FliG (17). During this process, FlhF enhances MS-ring formation by binding to the N-terminal region of FliF (18). A previous structural analysis of the C-ring in *Vibrio* using cryo-electron tomography revealed that the C-ring exhibits a 34-fold rotational symmetry, indicating that the *Vibrio* C-ring comprises 34 molecules of FliG (19). Since FliG and FliF interact in a one-to-one manner, it is assumed that the MS-ring also consists of 34 molecules of FliF in the native flagellar motor in *Vibrio*.

The structure of the *Salmonella* export gate complex—embedded in the M-ring center—was determined to be the FliP5FliQ4FliR1 complex (20). The MS-ring structure with the export apparatus and rod complex, the FliE6FlgB5FlgC6FlgF5FlgG24 complex, was determined in the native basal body of the *Salmonella* flagellum (21, 22). The flexible loops at the top of the S-ring interact with FlgB, FlgC, and FlgF, the inner surface of the S-ring interacts with FliE and FlgB, and the inner surface of the RBM2 ring interacts with FliQ and FliP to stabilize the export gate complex in the MS-ring (8, 21, 22).

The *fliE*, *fliF*, *fliG*, *fliH*, *fliI* and *fliJ* genes form operons on the chromosome of *Vibrio*. Based on the report of the functional FliF-FliG fusion protein in *Salmonella* (23), we cloned the *fliF* and *fliG* genes from *Vibrio* into the plasmid and modified them by deletion of a single nucleotide just 5’-end of the *fliG* start codon to overexpress *Vibrio* FliF-FliG fusion proteins (named FliFG fusion proteins) in *E. coli* (Fig. S1A) (24). The molecular weights of FliF and FliG are ∼64 and 39 kDa, respectively, and that of the FliFG fusion protein with an N-terminal His-tag is ∼104 kDa. *Vibrio* FliFG fusion proteins efficiently form the MS-ring in *E. coli* (24).

Here, we introduced FliG-G214S or FliG-G215A mutations into *Vibrio* FliFG fusion proteins. The FliG-G214S and FliG-G215A mutations confer strong switch bias and CW-locked rotation of the flagellar motor, respectively (25). The diameter of the C-ring top is 46.2 nm for the FliG-G214S mutant and 49.0 nm for the FliG-G215A mutant, induced by the FliG conformational change by the mutation (19). We performed single-particle cryo-EM of the purified MS-ring formed by FliFG fusion proteins with the FliG-G214S mutation and determined the near-atomic resolution structure of the *Vibrio* S-ring.

## RESULTS

### Purification of FliFG fusion protein with FliG mutation

We overproduced *Vibrio* His-FliFG fusion proteins with FliG-G214S or FliG-G215A mutation in *E. coli* cells. The membrane fraction containing the MS-ring formed by His-FliFG fusion proteins was treated with the detergent lauryl maltose neopentyl glycol (LMNG), and the MS-ring was precipitated by ultracentrifugation from the detergent-solubilized membrane fraction. We then tried to purify the MS-ring with the His-tag by cobalt-affinity chromatography; however, the MS-ring did not strongly bind to the cobalt column, and most of the MS-ring passed through the column due to unknown causes. The flow-through fraction was ultracentrifuged to precipitate the MS-ring, and the precipitated fraction was subjected to a size-exclusion column (Fig. S1B and C). The MS-ring was recovered near the void fractions as a very large molecular-weight complex. When the void fractions were subjected to SDS-PAGE, the FliFG fusion protein (∼100 kDa protein band) was detected (Fig. S1D and E). These fractions were observed using electron microscopy (EM).

### Cryo-EM structural analysis of the MS-ring formed by FliFG fusion proteins

We observed purified MS-rings formed by FliFG fusion proteins with FliG-G214S or FliG-G215A mutation using EM following negative staining (Fig. S2A and B). After the samples were concentrated using an Amicon device, they were applied to an EM grid and rapidly frozen for cryo-EM observation. The MS-ring structure with the FliG-G214S mutation was observed by cryo-EM (Fig. S2C); however, the MS-ring with the FliG-G215A mutation could not be observed, likely due to aggregation during the concentration process of the FliFG-G215A mutant proteins. Therefore, we performed single-particle cryo-EM of the MS-ring with the FliG-G214S mutation for structure determination. A total of 228,461 particle images extracted from 5,962 cryo-EM micrographs were analyzed. They were separated into two classes that showed the 35- or 34-fold rotational symmetry on the S-ring (Fig. 1C and D; Fig. S3). No other symmetry in the 2D class averages was observed, indicating that the purified MS-ring consisted of 35 or 34 subunits of the FliFG fusion proteins, and there was no other variation. Because the MS-ring in the native flagellar motor is thought to comprise 34 FliF subunits (19), it is inferred that the 34-fold rotational symmetry structure reflects the native S-ring structure. After postprocess without symmetry correction, their resolutions of 35 and 34 subunits MS-ring were 3.76 and 4.28 Å, respectively. Subsequently, we built the atomic model of the S-ring part (RBM3) at 3.23 and 3.33 Å resolutions, from the cryo-EM density maps of both classes after C35 and C34 rotational symmetry corrections, respectively (Fig. 1E and F, Fig. S3). We built the structural model for the S-ring, but due to poor density, could not build the model for the M-ring. It is noteworthy that weak densities corresponding to RBM2 were visible, similar in size to the previously reported RBM2 structures at the cross-section from side-views (Fig. 1C, D, G, and H), although it was impossible to build the atomic model from our maps.

The S-ring is formed by residues 257–449 of *Vibrio* FliF (FliF257–449), which corresponds to the latter half of the periplasmic region (Fig. 1B). The overall monomeric structure of our *Vibrio* S-ring was similar to previously reported monomeric structures of *Salmonella* S-rings (Fig. 2). The FliF257–449 monomeric structure consists of two structural regions: a conserved globular domain with an αββαβ motif (α1/β1/β6/α2/β7; residues 257–299 and 392–449) which forms a core of RBM3 (RBM3core), and a collar-like region (β-collar; residues 300–388) containing two sets of antiparallel β sheets (β2/β5 and β3/β4) with an invisible region (residues 328–366) at the top of the β-collar (Fig. 2A and B). The monomeric structures in the S-ring part from the 35- and 34-mer structures are almost identical (RMSD = 0.286 Å) (Fig. S5A and B). The monomeric structures of the *Salmonella* S-ring RBM3core were also highly similar to those of *Vibrio*; the RMSDs were less than 1.3 Å (Fig. S5C, Table S3).

**FIG 2.**
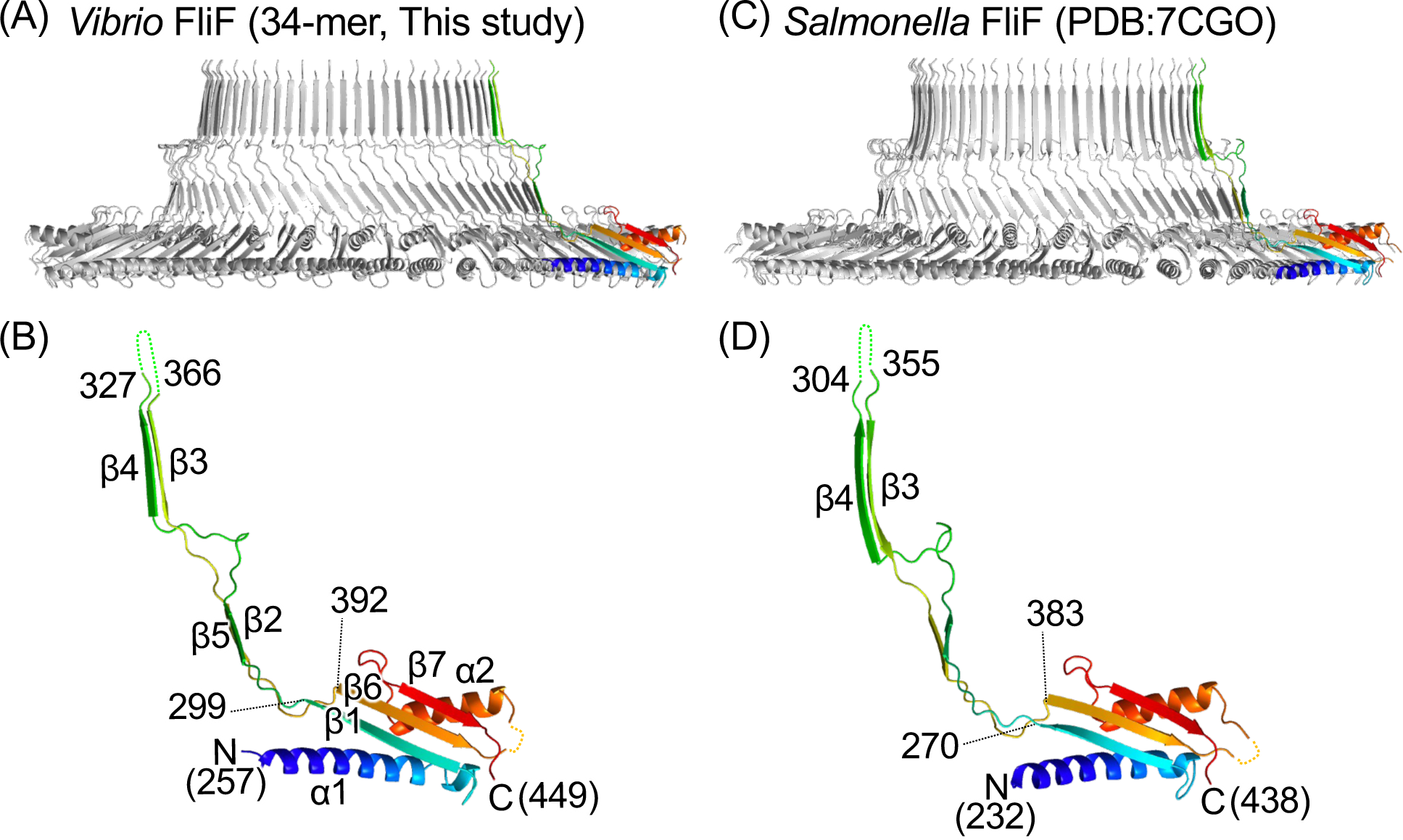
Comparison of the S-ring and FliF structures of *Vibrio* and *Salmonella*. (A) Cross-section view of the *Vibrio* S-ring structure shown as a Cα ribbon drawing. A protomer is colored in rainbow gradient from blue at the N-terminus to red at the C-terminus. (B) Magnified view of the protomer in (A). (C) Cross-section view of the *Salmonella* S-ring structure. (D) Magnified view of the protomer in (C).

### S-ring structure comparisons from *Vibrio* and *Salmonella*

We compared the *Vibrio* S-ring FliF 34-mer structure with three previously reported *Salmonella* S-ring FliF 34-mer structures (Fig. 3) where a notable difference is the tilt angle of the RBM3core domain. When measuring the α1 helix angle from the side-view of the ring, the RBM3core outer part is tilted downwards by 8° in the *Vibrio* S-ring, whereas it is tilted upwards by 6 or 8° or is nearly horizontal in the *Salmonella* S-ring (Fig. 3B). When comparing the β2/β5 sheet angles, they all exhibited a similar angle (55°) when viewed from the outside of the ring, whereas the angle is smallest in the *Vibrio* S-ring (58°) compared with the *Salmonella* S-ring (64–68°) when viewed from the side (Fig. 3B). The relative angle variation between the RBM3core and the β2/β5 sheet from the side-view indicates that the linker connecting them is structurally flexible. As for the β3/β4 sheet, all exhibit similar angles of ∼85–89° when viewed from the side (Fig. 3B). Residues 309–318 connecting β2 and β3 form a protruding triangular loop (β2-β3 loop) in the *Vibrio* S-ring, which is shorter than the loop (281–294) in the *Salmonella* S-ring (Fig. 3C and D, S6A). The relative angles between the β2/β5 sheet and the β3/β4 sheet are observed to be wider in the *Vibrio* S-ring (28°) compared with the *Salmonella* S-rings (19–25°) (Fig. 3B). This difference arises from variances in the shape of the β2-β3 loop in *Vibrio* and *Salmonella*.

**FIG 3.**
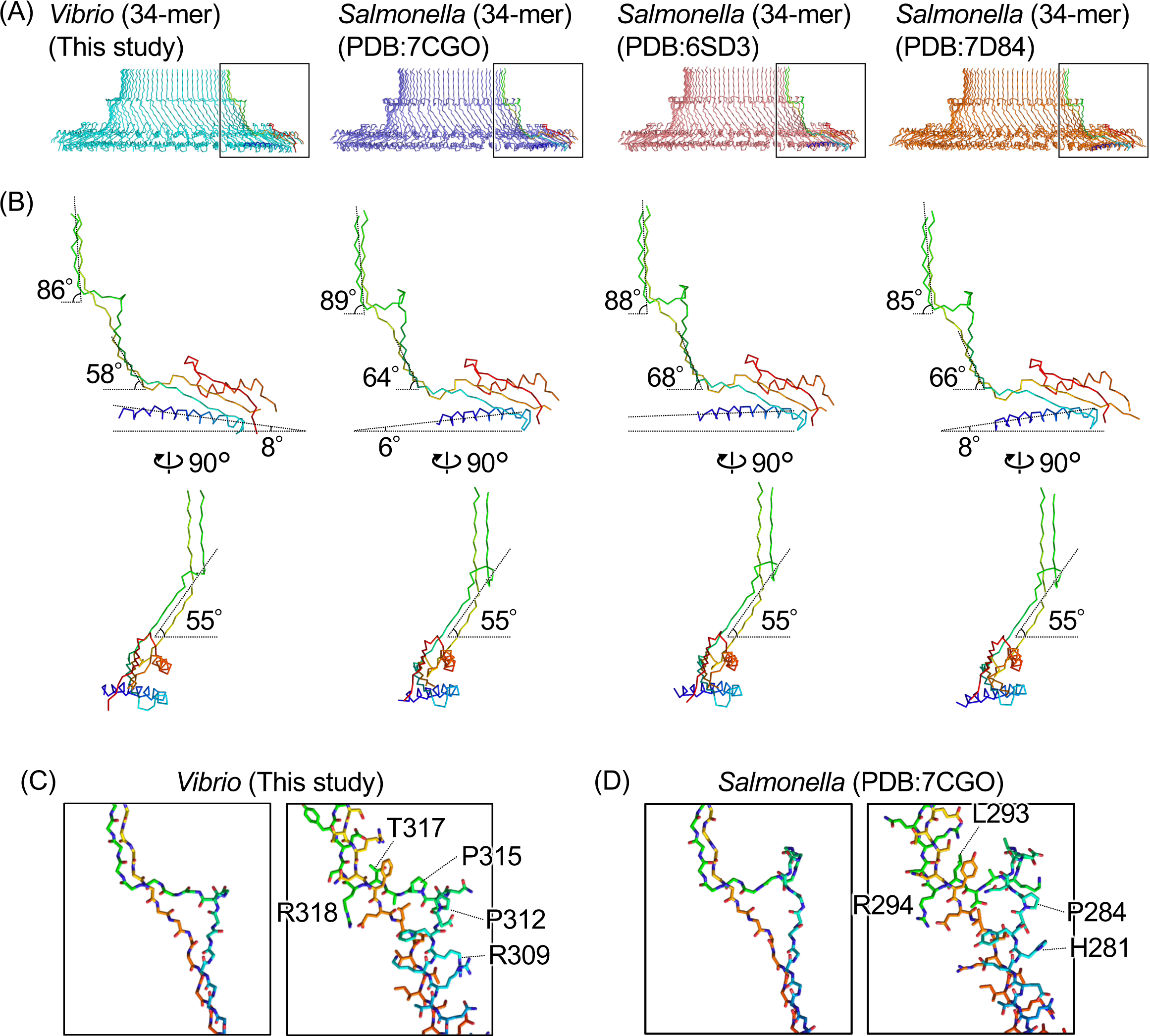
Comparison of the various S-ring and FliF structures. (A) Cross-section view of the Cα ribbon drawing of the S-ring structures from the MS-ring formed by *Vibrio* FliFG fusion protein (34-mer, this study), the intact flagellar hook basal body in *Salmonella* (PDB ID: 7CGO), the MS-ring formed by *Salmonella* FliF (PDB ID: 6SD3), and the MS-ring formed by *Salmonella* FliF (PDB ID: 7D84). All have 34-fold rotational symmetry. A protomer in each ring is colored in rainbow. (B) Comparison FliF protomer structures in the S-rings. The protomers colored in rainbow in (A) are shown. The inclination angles of the RBM3core, the antiparallel β2/β5 strands or the β3/β4 strands relative to the horizontal are shown. (C and D) Comparison of the protruding triangular β2–β3 loops in *Vibrio* FliF (this study) and *Salmonella* FliF (PDB ID: 7D84) structures. Left: stick models of the main chains. Right: stick models with the side chains.

When focusing on the subunit interface of the RBM3core during ring formation, interaction was facilitated by two sites: an electrostatic interaction between E270 and K276 and a hydrophobic interaction between A399, V429, and L447 in the *Vibrio* S-ring (Fig. 4A and B). These two interaction sites are conserved in the *Salmonella* S-ring, with an electrostatic interaction between E242 and R248 and hydrophobic interactions among V390, L415, and V433. In contrast, the *Salmonella* S-ring has additional hydrophobic interactions among L235, F237, and V241, which are not conserved in *Vibrio* (Fig. 4C and D), indicating that the interaction between the adjacent RBM3core regions in the S-ring is weaker in *Vibrio* than in *Salmonella*.

**FIG 4.**
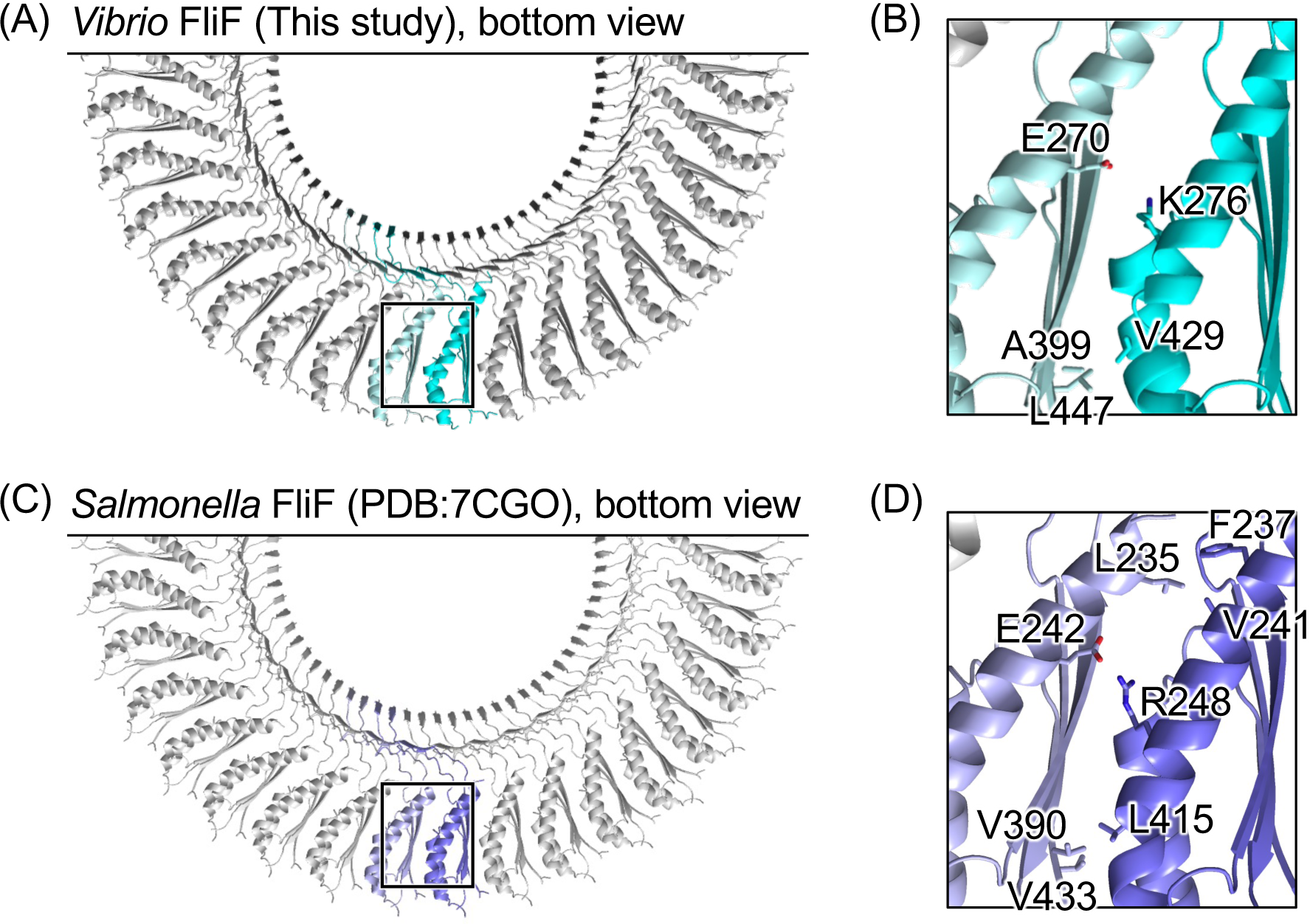
Comparison of the inter-subunit interface in the S-ring of *Vibrio* and *Salmonella*. (A and C) Bottom view of the Cα ribbon drawing of the S-ring structures of *Vibrio* and *Salmonella*. (B and D) Magnified view of the squared region in (A) and (C). The side chains contributing the inter-subunit interaction in the RBM3core are shown in stick models.

The internal surface of the S-ring is known to interact with the rod and export gate. Here we found the inner surface electrostatic distributions of the S-ring are different between *Vibrio* and *Salmonella* (Fig. 5A and B).

**FIG 5.**
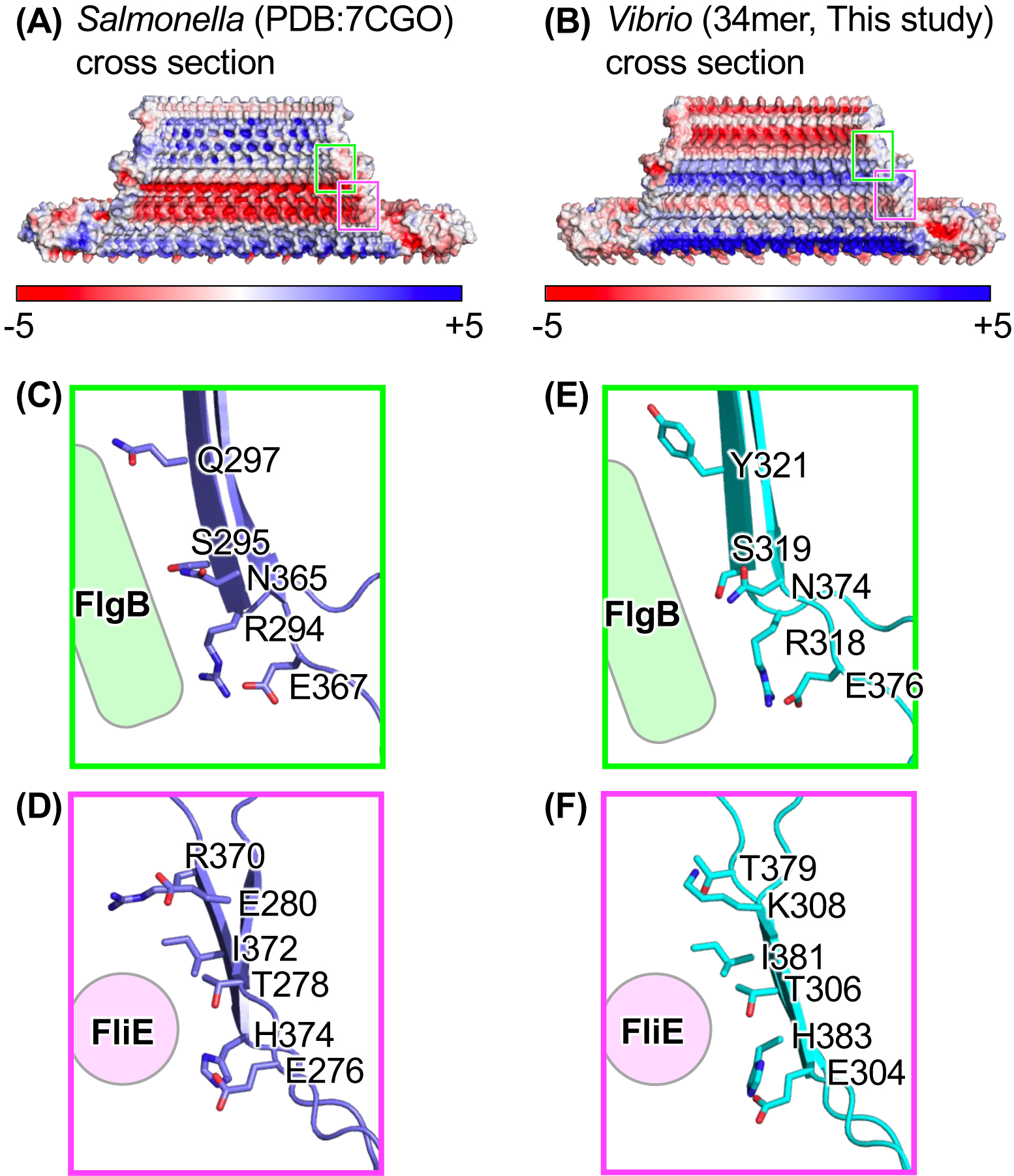
Electrostatic surfaces and regions interacting with rod proteins. (A and B) Cross-section image of the surface model with electrostatic potential of the S-ring structure. (C and D) Magnified views of the cyan and magenta squares in (A). (E and F) Magnified views of the cyan and magenta squares in (B). Sidechains contributing to the interaction with FlgB or FliE are shown in stick models.

In *Salmonella*, the S-ring internal surface directly interacts with specific regions of FliE and FlgB. Some residues at the inner upper part of the β-collar in FliF (R294/S295/Q297/N365/E367) interact with a flexible loop in the DC domain in FlgB (residues 58–82), and some residues around the base of the β-collar in FliF (E276/T278/E280/R370/I372/H374) interact with the N-terminal α-helix in FliE (Fig. 5C and D, S6 and S7) (21, 22, 26, 27). Most of the S-ring residues that interact with FliE or FlgB were conserved in our *Vibrio* S-ring structure, except for Q297 in *Salmonella* (Y321 in *Vibrio*) and E280 and R370 in *Salmonella* (K308 and T379 in *Vibrio*) (Fig. 5D and E, S6A).

## DISCUSSION

In this study, we determined the near-atomic resolution structure of the S-ring (RBM3) and the low-resolution structure of the RBM2 region in the MS-ring, composed of FliFG fusion proteins from *Vibrio*. We did not observe other regions, including RBM1, the two TM helices, C-terminal region of FliF, or fused FliG at the C-terminus of the FliFG fusion protein. Structural analysis of the purified MS-ring formed by FliFG fusion proteins using high-speed atomic force microscopy (HS-AFM) showed that it has a flexible structural region derived from fused FliG around the MS-ring structure (24). Here, we observed MS-rings consisting of FliFG fusion proteins with the FliG-G214S or FliG-G215A mutation by HS-AFM (Movie S1 and S2) and found vague structures at the outer part of the MS-ring, similar to the ring without the FliG mutation (24).

Our S-ring structures are composed of 34 or 35 molecules of the FliFG fusion protein, and the 34-mer structure should reflect the native S-ring structure, since the in situ MS-ring is a 34-mer of FliF (19). Although *Salmonella* FliF molecules can form MS-rings simply by overexpression alone in *E. coli*, *Vibrio* FliF molecules cannot form these by overexpression alone in *E. coli* but can by co-expression with FliG or FlhF or by fusion with FliG (24). The differences between *Salmonella* and *Vibrio* may be due to FliF characteristics. In fact, focusing on the inter-subunit interface of the RBM3core regions in the S-ring, *Vibrio* FliF exhibited less hydrophobic interaction than *Salmonella* FliF (Fig. 4). This difference would potentially elucidate why MS-ring formation by FliF alone is less likely to occur in *Vibrio*.

We also found that the tilt angle of the RBM3core in the *Vibrio* S-ring was different from that in *Salmonella* S-rings (Fig. 3B). Because SpoIIIAG—a 30-mer ring component in the feeding tube apparatus in *Bacillus subtilis—*also contains a RBM3core-like structure and the corresponding α1 helix is tilted upwards by about 23° (28), the tilt angle of RBM3core might have little effect on ring formation. The difference in relative angle between the β2/β5 and the β3/β4 sheets in the *Vibrio* S-ring and the *Salmonella* S-rings (Fig. 3B) is thought to arise from shape differences of the protruding triangular β2-β3 loop, with the length of the loop in *Vibrio* being four amino acids shorter than that in *Salmonella* (Fig. 3C and D). Due to such distinctive differences in the protruding triangular β2-β3 loop, it is presumed that the β3/β4 sheet is arranged vertically in *Vibrio* as well, regardless of tilt angle differences in the RBM3core.

The internal surface of the S-ring interacts with the FliE and FlgB rod proteins, and the FliF key residues for the interaction are well-conserved in *Vibrio* and *Salmonella*, except for K308, Y321, and T379 in *Vibrio* which correspond to E280, Q297, and R370 in *Salmonella* (Fig. 5C and D). It is possible that these non-conserved residues do not strongly contribute to the interactions with FliE or FlgB, as they are located slightly away from the interacting conserved residues. Alternatively, the non-conserved residues may participate in the interaction specific to *Vibrio*; in fact, the amino acid sequences of the DC domain in FlgB and the N-terminal α-helix in FliE—potentially interacting with the S-ring—are not fully conserved between *Salmonella* and *Vibrio* (Fig. S6B and C). Despite notable differences in the electrostatic distribution on the S-ring inner wall (Fig. 5A and B), key residues interacting with FlgB and FliE were conserved, indicating that the S-ring does not interact with the rod complex through its entire inner surface, but the specific residues on the inner wall are sufficient for the interaction between the S-ring and the rod.

## MATERIALS AND METHODS

### Strains, plasmids, and media

The bacterial strains and plasmids used here are listed in Table S1. *Escherichia coli* cells were cultured in LB medium [1 % (w/v) bactotryptone, 0.5 % (w/v) yeast extract, and 0.5 % (w/v) NaCl]. Ampicillin was added at a final concentration of 100 μg/mL. Point mutations (FliG-G214S or FliG-G215A) in plasmid pRO301 were introduced using the QuikChange site-directed mutagenesis method (Agilent).

### Purification of the MS-ring composed of the FliFG fusion proteins

An over-night culture of *E. coli* cells containing plasmid (pRO302 or pRO303) was inoculated at 1/50 dilution into 50 mL of LB medium containing ampicillin and grown at 37 °C for 4 h. The 20 mL culture was inoculated into 2 L of LB medium containing ampicillin and grown at 37 °C for 3–4 h. When optical density at 660 nm reached 0.4–0.5, isopropyl-β-D-thiogalactopyranoside was added at 0.5 mM, the culture was cooled on ice for 20 min, and then incubated at 16 °C for 20 h. The cells were collected by low-speed centrifugation and the pellet was stored at −20 °C if necessary. The pellet was resuspended in 25 mL of TK buffer (20 mM Tris-HCl [pH 8.0], 200 mM KCl) or TN buffer (20 mM Tris-HCl [pH 8.0], 200 mM NaCl) containing 1 mM EDTA.

The suspension was placed in a conical tube and sonicated three times at a power of 6 and 50 % duty ratio for 1 min. After low-speed centrifugation, the supernatant was recovered, and the precipitated cells were suspended in 25 mL of TK or TN buffer containing 1 mM EDTA and sonicated again under the same conditions as before (repeated twice in total). A 1/500 volume of 1 M MgCl2 was added to the pooled supernatants and centrifuged at low speed. The resulting supernatant was ultracentrifuged at 150,000 × *g* for 60 min. The precipitates were suspended in 20 mL of TK or TN buffer, and 2 mL of 10 % (w/v) LMNG was added and incubated at 37 °C for 30 min. After low-speed centrifugation, the supernatant was ultracentrifuged at 150,000 × *g* for 60 min. The precipitate was suspended in 1 mL of TK or TN buffer containing 0.01% (w/v) LMNG. The suspension was applied to a cobalt column, and the flow-through fraction was ultracentrifuged and suspended in 1 mL of TK or TN buffer containing 0.01 % (w/v) LMNG. The suspended solutions were subjected to the size-exclusion column (Superose 6 10/300, GE healthcare) equilibrated with TK100L buffer (20 mM Tris-HCl [pH 8.0], 100 mM KCl, 0.0025 % (w/v) LNMG) or TN100L buffer (20 mM Tris-HCl [pH 8.0], 100 mM NaCl, 0.0025 % (w/v) LNMG).

### Sample preparation and data correction of negative staining images by EM

Elution fractions containing the MS-ring composed of FliFG fusion proteins with the FliG-G214S or FliG-G215A mutations were concentrated 5-fold using an Amicon Ultra 100 K device (Merck Millipore). The G214S and G215A mutants were diluted 50- and 100-fold, respectively, in TN100L buffer. A 5 μL solution was applied to a glow-discharged continuous carbon grid. Excess solution was removed using filter paper, and the sample was subsequently stained on a carbon grid with 2 % (w/v) ammonium molybdate. Images were recorded using a H-7650 transmission electron microscope (Hitachi) operated at 80 kV and equipped with a FastScan-F114 CCD camera (TVIPS) at a nominal magnification of 40,000×.

### Cryo-EM observation

A concentrated MS-ring sample composed of the FliFG fusion protein with the FliG-G214S mutation was applied to a Quantifoil holey carbon grid (R1.2/1.3 Cu 300 mesh, Quantifoil Micro Tools GmbH) with glow-discharge treatment on one side of the grid. The grids were placed in liquid ethane cooled with liquid nitrogen for rapid freezing using a Vitrobot Mark IV (Thermo Fisher Scientific) with a blotting time of 7 s at 4 °C and 100 % humidity. The data were collected on Titan Krios electron microscope (FEI) equipped with a thermal field-emission electron gun operated at 300 kV and an Ω-type energy filter with a 20 eV-slit width. Image data sets of 6,372 micrographs collected by a Titan Krios microscope were automatically recorded on a K3 direct electron detector camera (Gatan) at a nominal magnification of 64,000 × corresponding to a pixel size of 1.14 Å with a defocus range from −0.7 to −1.7 μm, using the SerialEM software. Micrographs were taken after a total exposure time of 7.329 s, and an electron dose of 0.78 electrons/Å^2^ per frame. The 64 micrograph frames were recorded at a rate of 0.115 s/frame. The data collection and image analysis are presented in Fig. S3 and Table S2.

### Data Processing

A total of 6,372 micrographs were motion corrected, and their CTF values were estimated using the CryoSPARC package. A total of 500 micrographs were used for the initial particle picking using a blob picker. A total of 125,759 particles were automatically extracted from the micrographs, and 2D classification was performed three times to remove false particles. In total, 15,833 particles selected from the 2D classification were used as input templates for the template picker. A total of 2,059,366 particles were automatically extracted from the micrographs, and 2D classification was performed once to remove false particles, of which 860,145 were selected. Ab-initio reconstruction was performed to generate two initial 3D models using 512,737 particles selected from the 2D classification. Heterogeneous refinement with C35 symmetry was then performed using two of the three initial models to generate a 3D model with C35 symmetry. Heterogenous refinement was performed with 860,145 particles using two of three 3D models with C35 symmetry and 2D classification to remove false particles. The remaining 510,049 particles underwent heterogeneous refinement applied individually with C1, C34, and C35 symmetries to generate 3D models. The remaining 320,933 particles from heterogeneous refinement without symmetry (C1) underwent further heterogeneous refinement using the C34 and C35 models. Of these, 261,524 and 55,606 particles remained in classes with C35 and C34 symmetries, respectively. The particles belonging to each class underwent homogenous refinement with C35 and C34 symmetry to generate models for the final heterogeneous refinement to remove false particles. The remaining 183,915 and 43,546 particles with C35 and C34 symmetries, respectively, underwent homogeneous symmetry refinement and Global CTF refinement. Subsequently, the particles with C35 symmetry underwent homogenous refinement and a map was obtained at 3.23 Å resolution (Class 1; C35). To relax the symmetry, local refinement was performed and a map obtained at 3.76 Å resolution (Class 1; C1). Particles with C34 symmetry underwent homogeneous refinement and a map obtained at 3.33 Å resolution (Class 2; C34). Moreover, homogenous refinement without symmetry and homogenous refinement were performed and a map obtained at 4.28 Å resolution (Class 2; C1). The local resolution of the 3D volumes with C34 and C35 symmetries was estimated using the CryoSPARC package at 0.143 of its Fourier shell correlation threshold. The cryo-EM maps were deposited in the Electron Microscopy Data Bank under accession codes EMD-39763, EMD-39765, EMD-39761, and EMD-39764.

### Model building

The atomic model of the S-ring was constructed from the maps of Class 1 (C35) and Class 2 (C34) using *Coot* (29) and refined using *Phenix* (30). A summary of the model refinement is presented in Table S2. Structural comparisons and analyses were performed using PyMOL (Schrödinger) and Chimera (31). The atomic coordinates were deposited in the Protein Data Bank (www.pdb.org) under the accession codes 8Z4G and 8Z4D.

### HS-AFM observation and image analysis

HS-AFM imaging was performed using a laboratory-built HS-AFM operated in the tapping mode, as previously described (24). The MS-rings composed of FliFG fusion proteins were deposited on a bare mica substrate for HS-AFM imaging. After 5 min of incubation, the residual proteins were washed off using observation TN100L buffer. HS-AFM imaging was performed in the sample solution at room temperature.

## ACKNOWLEDGEMENTS

This study was partially supported by JSPS KAKENHI Grant Numbers 21H00393, 21H01772, 22K18943 (to T.U.), and 20H03220 (to M.H.).

## Author contributions

N.T., T.N., and M.H. designed the study; N.T., T.N., M.H., M.K., T.M., T.U., H.K., T.M., and M.H. performed the experiments; J.K., T.K., S.K., N.T., K.I., and M.H. analyzed the data; N.T., K.I., and M.H. wrote the manuscript.

